# Revisiting the habitat selection of the Eurasian Woodcock in winter: insights from the Mediterranean region

**DOI:** 10.1101/2025.05.19.654177

**Authors:** Camille Beaumelle, Jessica Barbet, Aurélie Cuby, Marc Chautan, Fabrice Etienne, Michel Martel, Alix Du Roure, Remy Chabanne, Estelle Lauer, Kévin Le Rest

## Abstract

Habitat selection is a key mechanism that enables animals to optimize their fitness in response to varying environmental conditions. Differences in habitat selection between populations in different geographical areas may indicate behavioral adaptations to local environmental conditions. Understanding the adaptive potential of species across broad geographical ranges is of primary interest to anticipate possible changes in species behavior or distribution in the context of climate change.

In this study, we investigated the habitat selection and the daily movement patterns of the Eurasian Woodcock, *Scolopax rusticola*, a bird species that winters across widely varying climatic zones. We tracked 84 individuals wintering in the Mediterranean regions with GPS-VHF transmitters, where climate and habitat conditions differ significantly from the regions with Atlantic climate influence, which have formed the main background for the ecology and behavior of this species in winter. To assess how woodcocks responded to varying habitat and environmental conditions, we collected data across three geographical regions spanning a gradient of Mediterranean climatic influence — ranging from northern subareas with denser forest and deeper soil to southern subareas characterized by less productive forests, garrigues, and rocky soil. In the northern Mediterranean region, woodcocks visited open habitats at night less than 53% of the time and less than 40% of the time in the two other regions with a stronger Mediterranean climate influence. This behavior was much less frequent than reported in studies conducted in areas with Atlantic climate influence (>80%). Woodcocks also changed their day/night activity patterns, as illustrated by their daily movements. They increased their daytime movements (11 to 29% higher) and reduced their nocturnal movements (12 to 18% lower) in the two regions with the strongest Mediterranean climate influence. During the day, when birds used only forested areas, denser forests were preferred in all studied Mediterranean subareas. Birds used different forested habitats between subareas, especially at night. For example, denser but shorter vegetation and higher rock cover were more strongly used at night in southern subareas. These forested habitats contrasted sharply with those in areas with Atlantic climate influence, the latter being plots rich in humus and deep soils.

Our findings highlight that basic ecological knowledge of species can be biased towards those known in certain types of habitats. They also underscore the remarkable behavioral flexibility of woodcocks, highlighting their potential to adapt to global change. However, the occurrence of escape movements under the driest conditions suggest that this change in behavior and habitat selection may be an early warning sign of the effects of climate change on the wintering areas. Overall, our study emphasizes the need to study the ecology of species across diverse environmental conditions to better understand their habitat requirements and adaptive capacity.

## Introduction

Global change is causing species to respond rapidly to new environmental conditions and altered biotic interactions (Albaladejo-Robles et al., 2023; IPBES, 2019; Ma et al., 2024; Zurell, Graham, et al., 2018). While empirical niche models predicting future species distribution highlight the urgency of planning conservation and management strategies to protect biodiversity, they often fail to capture species’ vulnerability or their potential for adaptation to rapid changes (Bellard et al., 2012; Dawson et al., 2011). Understanding these processes is essential for developing effective conservation strategies (Bellard et al., 2012; Dawson et al., 2011).

The study of habitat selection - defined as a modification in the disproportionate use of resources or conditions by an animal (Mayor et al., 2009) - is crucial for understanding the spatial distribution and abundance of species. It also helps to identify priority habitats that should be preserved or managed to support the persistence of species in a region, providing practical and effective tools for conservation strategies. Habitat selection occurs at multiple scales, both temporally and spatially (Mayor et al., 2009), sometimes across large and heterogeneous distribution areas that do not necessarily provide optimal conditions for the species. Some geographical areas may also be remote or difficult to access, further complicating data collection. Consequently, habitat selection studies often rely on data from a small part of an animal’s range and cover only a limited period of the animal’s lifespan (Fraser et al., 2018; Shaw et al., 2025, but see Beltran et al., 2025). The insights gained from such studies and the management recommendations derived from them are therefore rarely transferable to other contexts within the species’ range (Beyer et al., 2010) or to other taxa (Yates et al., 2018). This underlines the need for broader, integrative studies that take into account the spatiotemporal variability of habitat selection (Yates et al., 2018).

Moreover, habitat selection is a complex process shaped by the interaction between habitat availability and the behavioral motivations of individuals to meet their biological needs (Beyer et al., 2010; Roever et al., 2014). For example, habitat selection may be influenced by factors such as the physiology of the individual (e.g., age, Skórka et al., 2016), access to specific resources (e.g., landfill availability for *Larus fuscus*, Langley et al., 2021) and predation risk (Wasserlauf et al., 2023). Such variation illustrates a behavioral flexibility in habitat selection, which is particularly critical for species occupying habitats that are spatially heterogeneous or/and subject to temporal changes (e.g., climate and land use changes), as optimal space occupancy decisions may vary according to these variable conditions (Mabille et al., 2012; Merkle et al., 2022; Pearse et al., 2024; Shahsavarzadeh et al., 2023; Verzuh et al., 2021).

Migratory birds may occupy a remarkable diversity of environmental niches throughout their annual cycle (Jackson et al., 2019; Ponti et al., 2020; Zurell, Gallien, et al., 2018), showing their behavioral flexibility in optimizing the use of available resources. Despite this potential adaptability, they often exhibit strong fidelity to their breeding, wintering and stopover sites, likely because such consistency offers advantages such as more efficiency of movement and foraging, as well as better predator avoidance (Piper, 2011). However, when these sites are degraded, migratory birds may prioritize fidelity over relocation to alternative areas, a behavior that can sometimes result in maladaptation (Merkle et al., 2022). This trade-off suggests that species with the highest site fidelity may be particularly vulnerable to the challenges posed by rapid global change (Chan et al., 2023; but see Blackburn & Cresswell, 2016; Wellbrock et al., 2017). To cope with these changes, behavioral compensation within the habitats used could be an important mechanism that enables highly site-faithful species to adapt to local habitat alterations (Merkle et al., 2022). For example, one potential form of behavioral compensation is the adjustment of habitat selection within the occupied site, but few studies have investigated the ability of species to employ these strategies in response to environmental change (Mallet et al., 2023). Such change in habitat selection strategy is one of the behavioral mechanisms that may enable species to cope with global changes and persist in their environment (e.g., Bailey et al., 2019).

The aim of this study is to determine whether a migratory bird occupying a broad geographical wintering range can adjust its habitat selection strategy in relation to an environmental and climatic gradient. Specifically, we evaluated the behavior and quantified the habitat selection of Eurasian Woodcocks wintering in areas with Mediterranean climate influence. The drought periods during winter are expected to become more frequent and longer in the Mediterranean region (Raymond et al., 2019). Despite this, the woodcocks’ ecology remains largely unknown in this wintering region. Indeed, most of studies were carried out in regions of Europe with Atlantic climate influence being not affected by drought periods in winter (Arizaga et al., 2015; Braña et al., 2010; Duriez, Ferrand, et al., 2005; Duriez, Fritz, Said, et al., 2005; Yves Ferrand & Gossmann, 2009; Guzmán et al., 2017). Typically, the Eurasian woodcock is described as a species that commutes between open nocturnal foraging sites, where it feeds mainly on earthworms, and diurnal forest sectors, where they rest in shelter.

However, in southern France, observations of woodcocks in open areas at night are less frequent, suggesting that individuals wintering in Mediterranean regions may exhibit different behavioral strategies. An important point of this study was thus to compare the results obtained from Mediterranean region with those from other regions. To investigate this, we used GPS-VHF devices and tracked the movements of 84 wintering woodcocks across three subareas with varying degrees of Mediterranean climate influence. First, we analyzed general movement patterns during wintering to assess woodcock behavioral variation across Mediterranean subareas, having contrasted type of habitat, and environmental conditions (soil wetness and temperature). At the landscape scale (i.e., habitat patches within the home range, Johnson (1980)), we examined whether woodcocks maintained the typical pattern of selecting forest habitats during the day and more open habitats at night, as observed in other wintering regions. At the local scale (i.e., specific resources within the habitat patches, Johnson (1980)), we analyzed the habitat structures within the forests selected by woodcocks to identify environmental features associated with higher woodcock presence in each subarea. Finally, we compared our findings with studies from other climatic regions.

## Material and methods

### Studied species

The Eurasian Woodcock is a forest dwelling, medium-sized bird species with a long bill and short legs. It is distributed across the entire Palearctic region (Gonçalves et al., 2019). Its overall population trend remains unknown. However, population declines have been reported in some parts of its breeding range, especially in Europe (BirdLife International 2021). While some individuals are resident in parts of west-central Europe, the majority migrate between wintering areas - in western and southern Europe and northern Africa - and breeding areas - in the central and northern part of Palearctic (Gonçalves et al., 2019). This species shows site fidelity to both wintering and breeding areas (Yves Ferrand & Gossmann, 2009; Tedeschi et al., 2020). During the wintering period, woodcocks are generally solitary and remain in forests during the day. At night, most birds leave the forest to feed in open habitats, mainly grasslands and crops (Duriez, Ferrand, et al., 2005; Duriez, Fritz, Said, et al., 2005; Hoodless & Hirons, 2007; O’Neill et al., 2025). Under mild conditions, a higher proportion of birds stay in forested areas at night (Duriez et al., 2005). In the Mediterranean part of their wintering range, woodcock behavior has not yet been thoroughly studied, but individuals have been observed less frequently in open habitats at night, suggesting a different behavior than in regions with Atlantic climate influence. Thus, we expected woodcocks to select forests with lower tree cover and sparser vegetation at night, as an alternative of using open habitat.

To our knowledge, there is no study on the Eurasian woodcock showing any effect of age or sex on forested habitat selection but such differences occurred for the American woodcock (*Scolopax minor,* Slezak et al., 2024).

Woodcocks feed primarily on earthworms in winter (Bende & László, 2022; Duriez, Ferrand, et al., 2005; Granval, 1986; Hoodless & Hirons, 2007). Earthworms account for about 85% of their energy intake, with millipedes being the secondary prey (Granval, 1986). But recently, another case study identified beetles and millipedes as the main prey items in an area with Mediterranean climate influence (Aradis et al., 2019). Soil conditions can impact prey availability, and during extreme climatic events, such as prolonged frost (see Péron et al., 2011), woodcocks may be forced to leave their wintering area. In Mediterranean wintering area, prolonged droughts could have effects on food availability comparable to those of frost.

### Study area

Three subareas were selected in distinct sectors of southern France with Mediterranean climate influence (Figure 1), each of which is characterized by vegetation adapted to dry summers and strong winds (called “le mistral” and “la tramontane”) but is influenced to varying degrees by the Mediterranean climate (Table 1).

**Figure 1:**
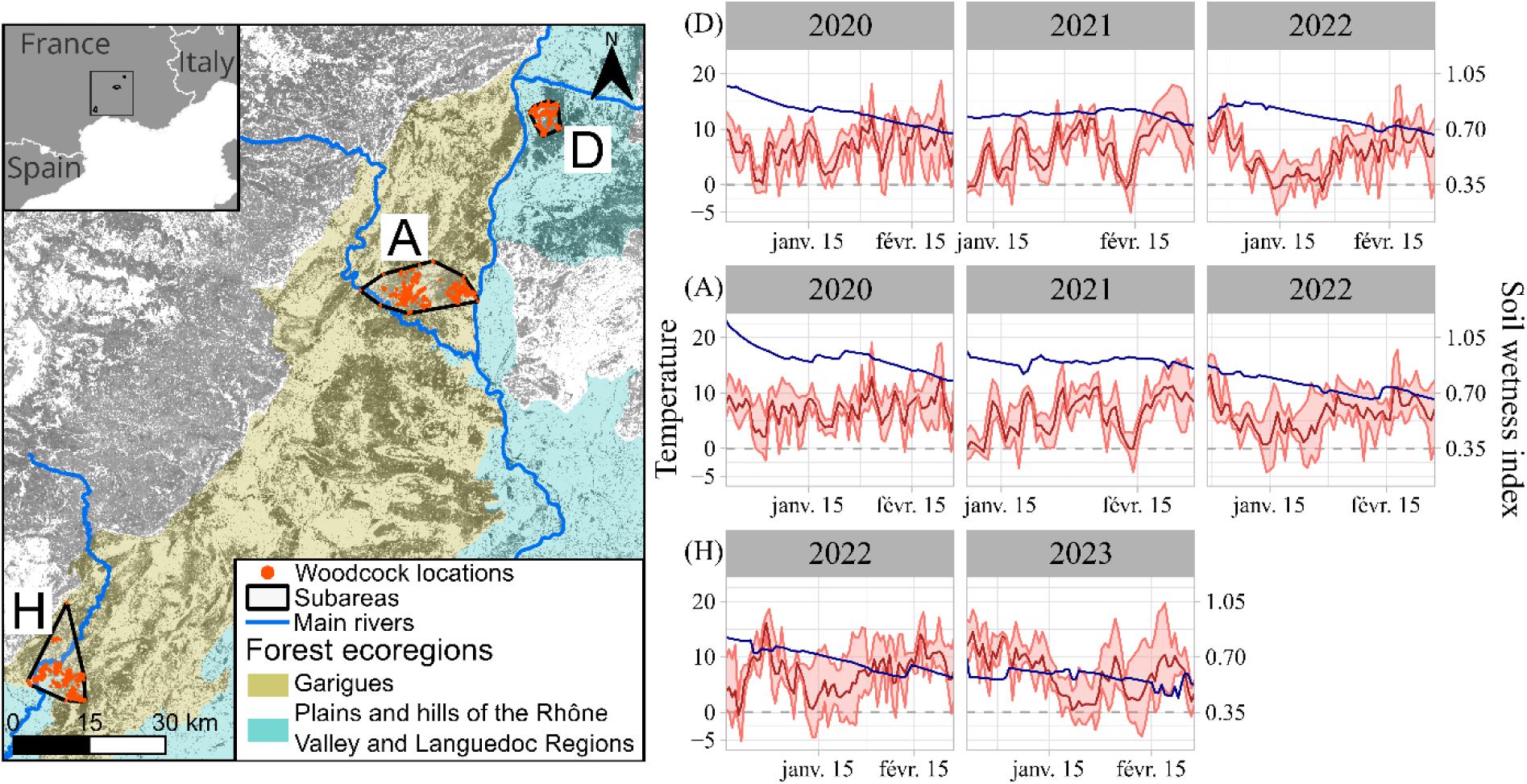
Location of the subareas and the recorded locations in southern France, as well as intra and interannual variation in temperature (red lines, daily min/max) and soil wetness (blue line). Subareas: D: Drôme, A: Ardèche gorges, H: Hérault gorges. The background of the map is the tree cover density, dark grey corresponding to high level of tree cover density (land.copernicus.eu/TCD2018).

**Table 1:** Descriptive data on the three subareas and sample size of woodcocks. N: number of woodcocks fitted with GPS, n: number of GPS locations per woodcock. Temperature and soil wetness were computed based on the data of the daily Safran model of Meteo-France (<u>data.gouv.fr/SIM</u>) during the tracking period. The surface area of subareas was calculated using the 100% minimum convex hull of woodcock locations.

| Subarea | Area (km<sup>2</sup>) | Year | N | Begin date | n<br>Mean [min-max] | Temperature<br>(°C)<br>Mean [min-max] | Soil wetness<br>Mean % [min-max] |
| --- | --- | --- | --- | --- | --- | --- | --- |
| (D) | 29 | 2019-2020 | 8 | 21<sup>st</sup> Dec. | 187 [61; 261] | 7 [-2; 19] | 80 [67; 97] |
|  |  | 2020-2021 | 6 | 9<sup>th</sup> Jan. | 126 [59; 170] | 7 [-5; 18] | 80 [72; 83] |
|  |  | 2021-2022 | 5 | 25<sup>th</sup> Dec. | 162 [115; 234] | 6 [-5; 18] | 76 [66; 85] |
| (A) | 150 | 2019-2020 | 7 | 21<sup>st</sup> Dec. | 168 [62; 273] | 7 [-3; 21] | 92 [76; 116] |
|  |  | 2020-2021 | 10 | 7<sup>th</sup> Jan. | 173 [144; 183] | 6 [-5; 17] | 91 [84; 97] |
|  |  | 2021-2022 | 10 | 29<sup>th</sup> Dec. | 143 [40; 224] | 6 [-5; 18] | 75 [63; 90] |
| (H) | 118 | 2021-2022 | 20 | 16<sup>th</sup> Dec. | 141 [42; 207] | 8 [-3; 18] | 59 [50; 67] |
|  |  | 2022-2023 | 18 | 21<sup>st</sup> Dec. | 182 [62; 257] | 8 [-4; 20] | 45 [40; 51] |

Subareas: D: Drôme, A: Ardèche gorges, H: Hérault gorges

The Mediterranean subarea of the Drôme (D) belongs to the ecoregion of the Plains and Hills of the Rhône Valley and Languedoc Region (see Figure 1, IGN, 2013). This subarea is relatively humid due to the influence of the continental and mountain climate, has moderately deep soils, and the forests are dominated by *Quercus pubescens* (a deciduous tree species). The Ardèche (A) and Hérault (H) gorges belong to the same forest ecoregion, the *garrigues* (or shrublands), as defined by the French National Geographic Institute (Figure 1, IGN, 2013). Their soils are shallower and much rockier than further north, so they drain faster, leading to drought conditions, mainly in summer but also during some winters (e.g. in 2022 and 2023). The *garrigues* are characterized by low, slow-growing vegetation. *Quercus ilex*, is the main tree species, which is evergreen. Others are *Pinus halepensis*, *Pinus pinea, Quercus coccifera*, *Erica* spp., *Cistus* spp. and *Arbutus unedo*.

The daily temperature fluctuated between −5°C and 20°C in the region, with few freezing days (Table 1, Figure 1). The soil wetness index (hereafter SWI) generally decreased significantly during the wintering period, e.g. from 0.97 to 0.68 in winter 2019-2020 (hereafter winter 2020) in D. The study period exhibited strong inter-annual variation in SWI: precipitation levels were high during the winter 2020, resulting in elevated SWI values in D and A at the beginning of the wintering period. Conversely, the winter 2021-2022 (hereafter winter 2022) was dry, leading to a drought period with SWI values <0.5 (Fiche Climat, 2015). The SWI in H was consistently lower than in the other subareas.

### Capture and tracking

The study was conducted from winters 2020 to 2022-23 (hereafter winter 2023). For each subarea, woodcocks were caught over two or three consecutive winters, from late December to early January (Table 1), i.e., a period where the individuals stayed on their wintering site (Ferrand et al., 2008), reducing the risk that individuals leave the studied subareas. The captures took place mostly at night in fields, vineyards or orchards, where the birds were detected and immobilized using a spotlight (dazzled) and then caught with a landing net (Gossmann et al., 1988). In the Hérault gorges, half of the birds were caught at night near ponds using vertical nets (mist-nets) set up before dusk, as woodcocks sometimes bathe for a few minutes in these ponds at sunset. The woodcocks were ringed and aged by plumage characteristics (Y. Ferrand & Gossmann, 2009). To determine sex, a feather was collected and analyzed by the CIBIO/CTM laboratory in Portugal (DAN analyses, Griffiths et al., 1998). If the individual weighed more than 300 g and was in good physical and health condition, it was equipped with a GPS-VHF tag (PinPoint VHF 240, Lotek Ltd, ∼8 g) using a leg-loop harness made with an elastic rubber cord (1.5 mm EPDM) lined within a silicone sheaths (IN 2.0; OUT 2.6 mm) to limit friction. Tag plus harness weighed a total of ∼ 10 g, representing less than 3% of bird’s body weight. The elastic rubber cord sometimes broke too early, i.e., a few weeks after a bird’s release, so we use a silicone tube instead of an elastic rubber in 2023 in H (harness being even lighter, ∼1g).

Each bird was first located by triangulation, and the data (GPS positions) were downloaded every two weeks, from approximately 50-100 meters to avoid any disturbance (depending on vegetation and topography). The tags were programmed to record one location every 6 hours, comprising two nocturnal locations (9 pm and 3 am) and two diurnal locations (9 am and 3 pm). In addition to the GPS data, the device recorded temperature, elevation, and GPS quality metrics (HDPO, eRes, number of satellites used for the location).

### GPS data filtering and selection

During the study period, we tagged 97 woodcocks. However, one bird was killed by a hunter a few days after being equipped, another was predated, and for six of them we could not successfully download the GPS data (the birds may have flown out of the study area or been predated, or the tags were defective). Five woodcocks wintering outside the three subareas were not considered in this study. Therefore, we analyzed the movements of 84 woodcocks here (Table 1). We carefully checked and filtered GPS locations used for analysis. All filtering steps and the corresponding sample size are provided in Appendix 1 (Figure A1).

Eight woodcocks in the Hérault gorges (subarea H) escaped during the driest period of the study and later returned to their primary wintering area. Data from these escape locations (18 to 74 km away) and points between the primary wintering area and the escape area were excluded from the analysis, resulting in the removal of 413 locations (24% of the total from these eight individuals).

As our study focused on the wintering period, we excluded locations recorded in March and April because woodcocks may change their habits to prepare for migration (Duriez, Fritz, Said, et al., 2005; Tedeschi et al., 2020).

### Statistical analysis of movement patterns

To analyze the global movement of woodcock, we computed diurnal and nocturnal distances as well as the distances between diurnal and nocturnal sites (commuting flights). All distances were calculated between pairs of consecutive locations using the “st_distance” function of the *sf* package (Pebesma et al., 2024). Commuting distances were estimated by measuring the distances between consecutive nocturnal and diurnal locations (see Duriez, Fritz, Said, et al., 2005). These included the distances of evening commute flights (occurring between 3 pm and 9 pm) and morning commute flights (occurring between 3 am and 9 am). Nocturnal and diurnal movements were estimated by measuring the distances between consecutive nocturnal locations (9 pm and 3 am) and between consecutive diurnal locations (9 am and 3 pm), respectively. We tested whether distances differed between woodcocks of different age and sex, and whether they varied according to subareas, temperature and soil wetness, by fitting a linear mixed model, using the *lme4* package (Bates et al., 2015). We included individuals as a random intercept to account for repeated measures and variability in behavior. The distances were log-transformed to achieve normality. Temporal autocorrelation was controlled by including Julian day as an explanatory variable. The daily soil wetness (SWI), the daily mean of temperature and the julian days were standardized. The three models (diurnal and nocturnal distances, and commuting flights) were fitted using the following formula, where subarea is a factor with three levels (D, A, H). The julian day covariate started on December 16^th^ for each year.

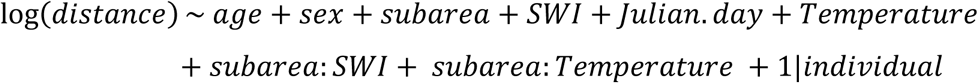

In addition, we analyzed the nocturnal commuting frequencies in open areas. To determine if a woodcock was either in a wooded or an open area at night, we used a high-resolution tree cover density (TCD) layer (10m raster from (<u>land.copernicus.eu/TCD2018</u>). The TCD for each location was calculated as the mean value of raster cells within a 30 m radius buffer to account for slight inaccuracies in GPS locations (9 ±10 m, estimated from data collected by transmitters left in the forest). Then, we defined a threshold based on the TCD values to distinguish strict open areas (TCD≤5%) from most forests, hedgerows, garrigues and orchards (TCD>5%). The response variable “open area”, (coded 0/1) indicated whether a woodcock was in an open area at night (1: TCD≤5%) or not (0 TCD>5%). We used the same explanatory variables as in the distance models. We fitted a binomial generalized mixed model as follows:

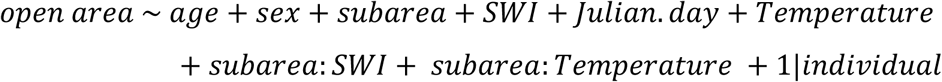

Confidence intervals were calculated using the *profile* method of the *confint* function of the *lme4* package (Bates et al., 2015). The p-values were produced by the Satterthwaite approximation with the *lmerTest* package (Kuznetsova et al., 2017), and the Nakagawa’s conditional R² (Nakagawa & Schielzeth, 2013) were computed with the *performance* package (Lüdecke et al., 2021). The diagnostic and the assumption of the model assumptions was checked with the DHARMa package (Hartig, 2024, Appendix 2, Figure A1).

### Statistical analysis of habitat selection at landscape scale

To analyze the habitat selection of woodcock at the landscape scale, we determined whether woodcock locations were situated in forested or more open habitats, using the continuous TCD values.

To measure the degree of TCD’s selection by woodcock, we fitted a resource selection function using logistic regression to model use-availability data (Boyce et al., 2002; Fieberg et al., 2021; Northrup et al., 2022). Habitats used by woodcock (based on GPS locations) were classified as 1, while randomly sampled available habitats were classified as 0 (hereafter named pseudo-absence). We added individuals random intercepts to account for variability in behavior between individuals as well as a different number of locations among individuals (Gillies et al., 2006). In our study, several woodcocks frequently commuted between areas more than 1 km apart, a behavior that was also observed in other wintering areas (e.g., in Brittany: Duriez, Fritz, Said, et al., 2005; in Spain: Guzmán et al., 2017) and some reach 5 km. Therefore, for each woodcock, the available habitat was defined as the area within a buffer of 1 km, 3 km or 5 km around all locations and we compared the results across the buffers considered. For each true location, 100 pseudo-absences were sampled, excluding areas never used by woodcocks, i.e., water bodies or artificial surfaces determined by the Corine Land Cover layer (100m raster from <u>copernicus.eu/clc2018</u>). Predictor variables included the TCD, the period of the day (day/night factor), the three subareas, the daily soil wetness (SWI), the daily mean temperature, the Julian date, the sex and the age. Continuous variables were standardized. The day/night factor for the pseudo-absence data was defined from the nearest available GPS location from each woodcock because some parts of the individual home range were more frequently used day time than at night and vice versa. The model was fitted using the following formula:

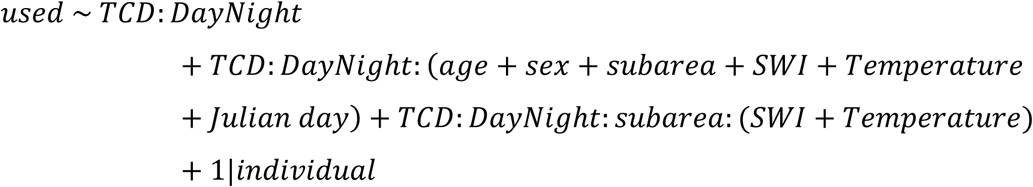

We repeated the random sampling of pseudo-absence locations 10 times for each buffer size (1, 3 and 5 km) to avoid results being too sensitive to one sampling, then averaged the 10 models using the default *model.avg* function of the *MuMIn* R package (Bartoń, 2024). the Nakagawa’s conditional R² (Nakagawa & Schielzeth, 2013) were computed with the *performance* package (Lüdecke et al., 2021). The diagnostic and the assumption of the model assumptions was checked with the DHARMa package (Hartig, 2024, Appendix 3, Figure A1).

To interpret the average effect of covariates on selection along the TCD gradient, we computed the relative selection strength (RSS, Avgar et al., 2017) associated with TCD-covariate interactions. For each covariate, RSS was calculated as exp(β_i_), where β_i_ is the coefficient of the interaction term between TCD, Day/Night, and the focal covariate in the fitted model. For interactions involving the subarea covariate, e.g., TCD x Day/Night x subarea x SWI, the average effect was calculated for a given subarea by combining the relevant interaction terms. For example, the RSS of TCD during the night in A as a function of SWI was computed as exp(β_TCD:night:A:SWI_) x exp(β_TCD:night:SWI_). RSS can be interpreted as the relative probability of use of two locations that differ by one standard deviation in a continuous covariate, or by a factor level for categorical covariates, while all other covariates are held constant (Fieberg et al., 2021). In our model An RSS value >1 indicates selection for locations with higher TCD values relative to the reference, whereas an RSS value <1 indicates selection for lower TCD values.

### Statistical analysis of forest habitat selection at local scale

To determine the environmental characteristics of forest habitats selected by woodcock at a local scale, we visited forest plots of 25 x 25m having one or more GPS location of tracked woodcocks (Appendix A4).

The habitat variables of interest chosen to describe these forested plots were based on the known ecology and behavior of the species. Woodcock is known to prefer young forests of small height (9-15 years), with moderate tree density, dense vegetation in the lower layers (herbaceous and shrub layers) but with clear areas to allow movement in the ground layer (Ferrand & Gossmann 2009). The variables measured characterizing the vegetation structure were thus: main tree species, dominant tree height, cover, proportion and distribution of grass, and the vegetation density at 10 cm, 50 cm, 100 cm and 150 cm (Table 2). In the studies subareas, rocks could completely cover the soil, potentially preventing woodcocks from probing the ground in search of soil invertebrates. Consequently, we also measured the proportion and distribution of rocks. After discarding or modifying redundant local habitat variables (tree species, distribution of rocks, and the proportion and distribution of grass, Appendix A4, Figure A3) and checked for correlations, we retained six habitat variables (Table 2) as explanatory variables for the model (Table 2). The maximum correlation calculated between these variables was 0.47.

**Table 2:** Description of the six retained habitat variables measured in plots located in forest.

| Habitat variables | Description |
| --- | --- |
| Dominant height of tree | Height $<4$ meters or $>4$ meters (binary) |
| Cover | Proportion of the ground covered by vegetation of $>0.5$ meters |
| Vegetation density 150 cm | Probability of not seeing the mark at 150cm |
| Vegetation density 50 cm | Probability of not seeing the mark at 50cm |
| Forest herb distribution | Scattered, connected covering <50% of the ground or<br>connected, covering ≥50% of the ground (3 levels<br>categorical factor) |
| Proportion of rocks | Proportion of the ground covered by rocks outcrops<br>and loose rocks |

To analyze forest habitat selection, we tested whether some plots were preferred by woodcocks, by quantifying the intensity of plot use, i.e., the number of nocturnal and diurnal locations per woodcock in the corresponding sampled plots during the studied period (Table 1). To account for overdispersion in the count data, we fitted a generalized linear model with a negative binomial distribution. A log-transformed offset in the model controlled for the different total number of locations per individual. During the day, woodcocks were always in forest; therefore, the offset for diurnal locations corresponded to the total number of diurnal locations. At night, woodcocks could use either forest or open areas. Nocturnal locations where the mean TCD exceeded 5% within a 12.5 m radius buffer (to match the 25 m x 25 m plot resolution) were considered as forest habitat.

We compared the performance of 6 competitive models composed of habitat variables (Table 2) and their interaction with age, sex, day/night, and the subareas. These included the null model, the model with habitat variables only, the model with habitat variables interacted with the period of the day (day/night), and three models in which the habitat variables also interacted with the period of the day, as well as with age, sex, or the subareas. Models were ranked based on their AICc values, and we selected the model with the lowest AICc value. Models with ΔAICc <2 were considered equivalent. In this case, we selected the most parsimonious model (Burnham & Anderson, 2004). The diagnostic and the assumption of the model assumptions was checked with the *DHARMa* package (Hartig, 2024, Appendix 4, Figure A4), the Nagelkerke’s R² (Nagelkerke, 1991) was computed with the *performance* package, and the estimated marginal means were obtained with the *emmeans* package (Lenth, 2025). Estimated marginal means represent model-based predictions evaluated for a focal variable within each subarea, while other continuous covariates are set to their average and factor effects are averaged over their levels.

We used R 4.4.1 (R Core Team, 2024) for data processing and statistics, and QGIS 3.28.1 (QGIS.org, 2024) and Inscape 1.3.2 (Inkscape Project, 2023) for data visualization.

## Results

In this study we analyzed the movements of 84 woodcocks, including 21 1^st^ winter male, 22 1^st^ winter female, 24 adult males and 17 adult females. From the start of the tracking record (Table 1) to the end of February, we collected a total of 13 482 locations with an average of 183 locations per woodcock (min= 45; max=280).

### Movement patterns

The movement patterns of woodcocks varied greatly depending on the individual and subarea (Appendix 2, Figure A2, Table A1). During the day, woodcocks almost always stayed in tree-covered areas (Figure 2). Only 1% of diurnal locations were recorded in open areas (TCD<5%). Most of these locations were in *garrigues* with sparse vegetation (68/87), and the rest (19/87) were in hedgerows, orchards, or crops. The use of nocturnal habitat at the landscape scale varied between individuals. There were 19 individuals that frequently used open habitats at night, with more than 50% of their nocturnal locations in non-tree-covered areas. In contrast, 65 individuals used open areas at night less than 50% of the time, including 27 woodcocks that had less than 5% of their locations in non-tree-covered areas.

**Figure 2:**
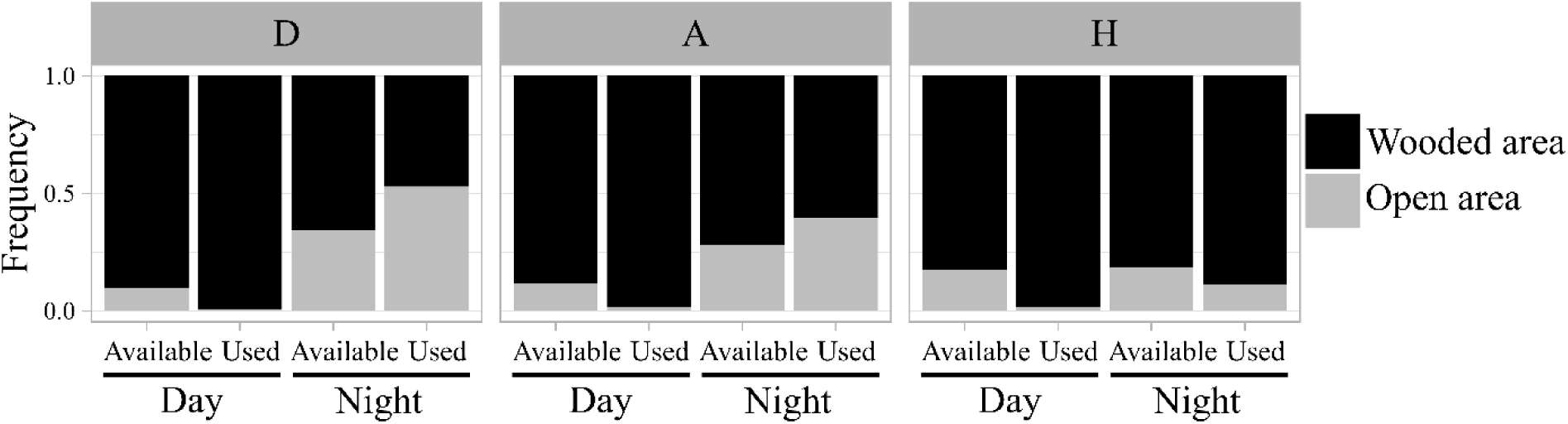
Frequency of open areas and wooded area in used habitats in comparison with the available habitats, computed with a buffer size of 3 km around used bird’s locations. The area was considered as forest if the tree cover density (TCD) was ≥ 5%. Each bar displays the mean habitat composition observed for all individuals pooled within the same subarea during nighttime and daytime. Subareas: D: Central Drôme, A: Ardèche gorges, H: Hérault gorges.

Sex and age had not significant effect on movement patterns (Table 3). Woodcocks in H commuted almost three times less frequently than those in D and nearly twice less frequently than those in A (Appendix 2, Table A2), and their commuting distances were shorter (Table 3). Over the course of the wintering period, woodcocks commuted slightly more often to open sectors at night (β_Julian day_ = 0.13 [0.03; 0.24]_CI95%_, Appendix 2, Table A2).

**Table 3:**
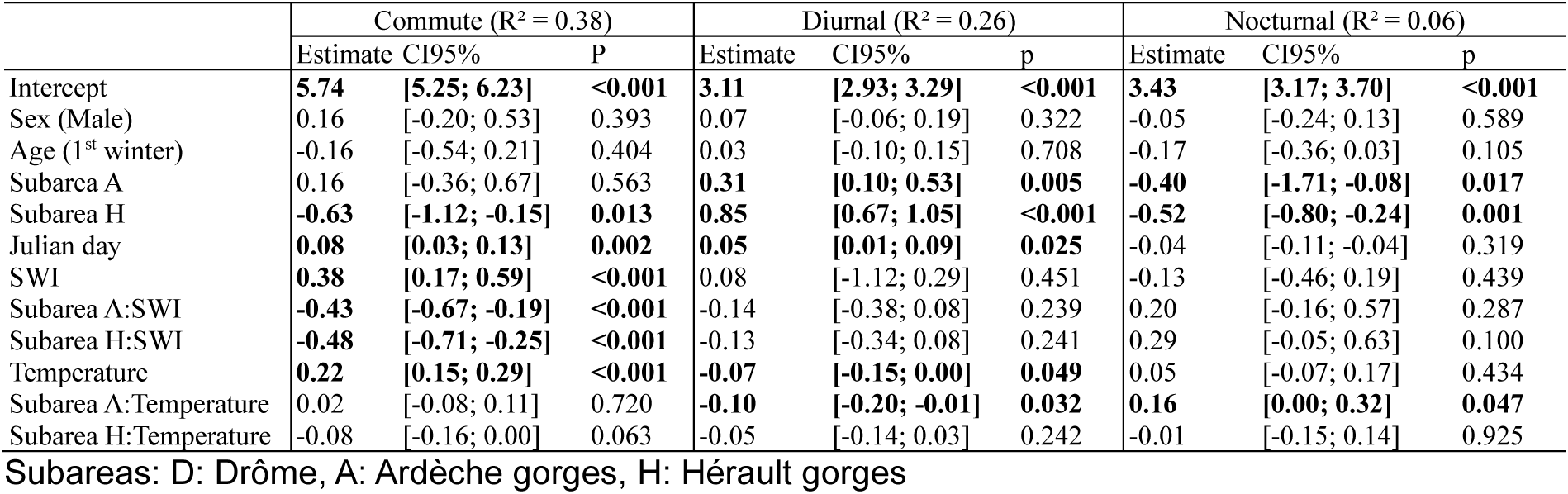
Estimates and 95% confidence intervals of the linear mixed models explaining commute, diurnal and nocturnal log-distances. Females and adults were the reference levels for the sex and age factors. The subarea D was the reference factor level. SWI: soil wetness index.

Woodcocks in A and H travelled more distance during the day than those in D (e.g., +11% in A and +29% in H relative to D, with T = 7°C and SWI = 0.7), but travelled less distances at night (e.g., −12% in A and -18% in H compared to D with T = 7°C and SWI = 0.7), as indicated by the diurnal and the nocturnal distance models (Table 3). During the day, distances were also slightly greater over the course of the wintering period (Table 3).

Under warmer temperatures, woodcocks commuted at longer distances (Table 3) and more frequently between forested diurnal and open nocturnal sectors, particularly in A and H. For instance, with SWI = 0.7 and T = 0°C, commuting frequency was 20% in D and 6% in A; at T=10°C, it doubled in D (41%) and quadrupled in A (26%). Diurnal distances decreased with increased temperature in all subareas (Table 3), while nocturnal distances increased with temperature in A only (Table 3).

Wetter conditions also increased the commuting frequency between forested diurnal and open nocturnal sectors across all subareas (β_SWI_= 0.94 [0.56; 1.33]_CI95%_, Appendix 2, Table A2). For instance, in A, at T = 10°C, commuting frequency rose from 26% at SWI=0.7 to 37% at SWI=0.8. In contrast, the effect of SWI on commuting distance differed among subareas: distances increased with SWI in D (e.g., at T=10°C, 310m at SWI=0.7 vs. 403m at SWI=0.8), but decreased with SWI in A and H (e.g.in A at T=10°C, 428m at SWI=0.7 vs.415m at SWI=0.8).

### Habitat selection at the landscape scale

Woodcocks that remained in the forest at night generally stayed in areas with lower TCD at night than during the day (Appendix A2, Figure A3).

Parameter estimates were generally consistent regardless of the size of the buffer used to sample the pseudo-absences around the bird’s location (Figure 3, Appendix 3, Table A1). Therefore, we primarily report the results from the 3 km buffer model (Nakagana’s R²= 0.07).

**Figure 3:**
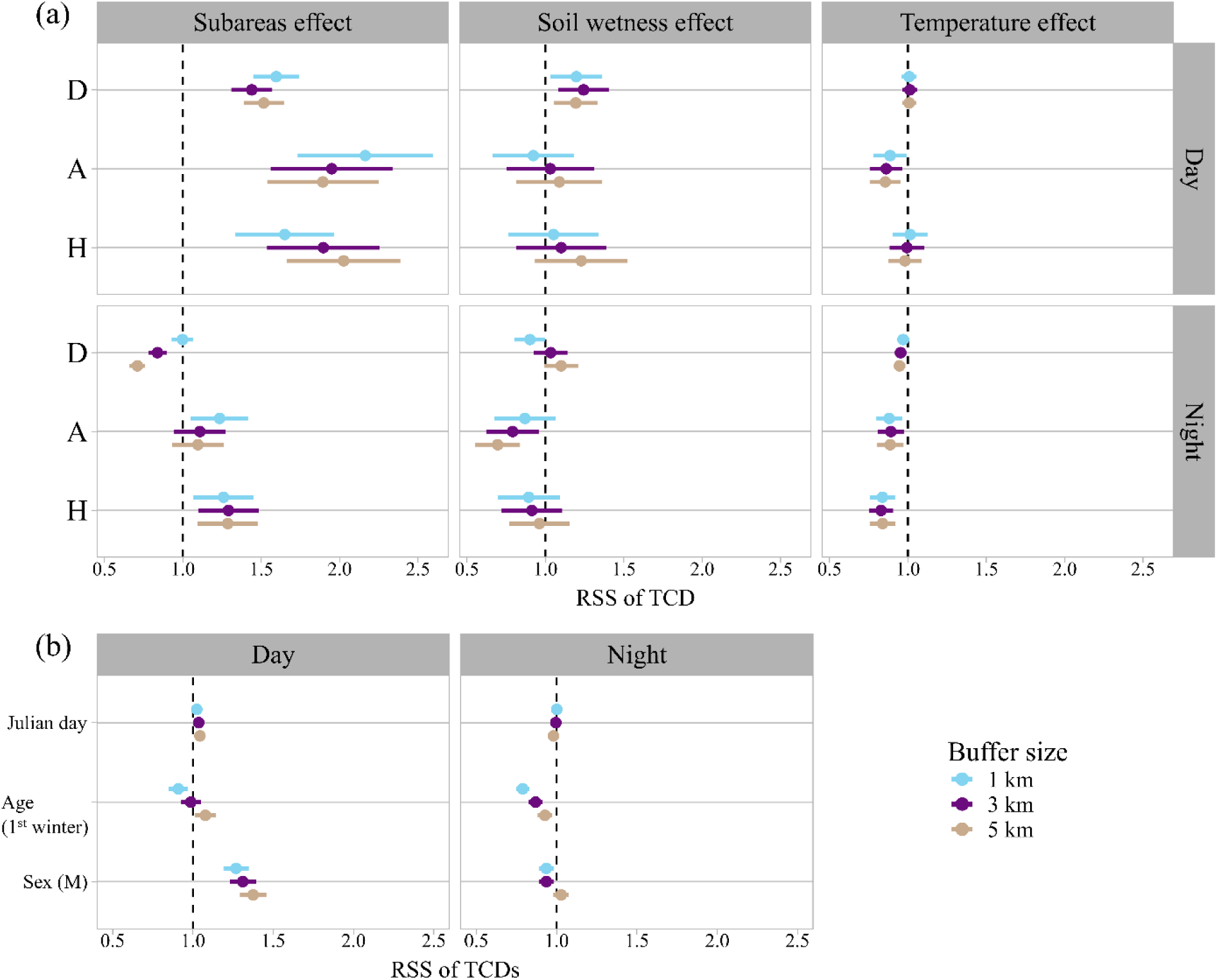
Relative selection strength (RSS) of the tree cover density (TCD) at the landscape scale obtained from the model averaging. (a) Specific selection depending on the subarea, and the soil wetness and the temperature in each subarea (b) Global selection depending on age, sex and Julian day in the three subareas. The available habitat was computed with a buffer size of 1 km, 3 km and 5 km. Females and adults were the references levels for the sex and age factors. Subareas: D (reference factor): Drôme, A: Ardèche gorges, H: Hérault gorges.

During the day, woodcocks significantly selected habitats with a higher tree cover density (TCD) than available in the surrounding area (e.g., in D: mean used TCD = 72 %, mean available TCD = 65 %, Figure 3). This selection was stronger in males (RSS = 1.31 [1.23; 1.39]_CI95%,_ Figure 3) and under wetter conditions in D (RSS = 1.23 [1.08; 1.40]_CI95%,_ Figure 3). Under milder temperatures, woodcocks in A selected lower TCD (RSS = 0.86 [0.76; 0.96]_CI95%,_ Figure 3).

At night, the results were more heterogenous between subareas. In D, woodcocks selected habitats with lower TCD when considering the 3- and 5- km buffer (RSS = 0.71-0.84, Figure 3) but this result was not significant when considering the 1km buffer. In contrast, TCD selection at night was slighly positive in A (only significant considering the 1km buffer) and positive in H (RSS=1.28 [1.13; 1.49]_CI95%_, Figure 3). Under drier conditions, selection for higher TCD was significant in A (RSS=0.77 [0.62; 0.96]_CI95%_, Figure 3). Woodcocks were also more likely to use less tree covered areas under milder temperatures, particularly in H (RSS=0.82 [0.75; 0.91]_CI95%_, Figure 3). 1^st^ winter birds showed stronger selection for lower TCD than adults at night (Figure 3). However, 1^st^ winter birds were not more likely to commute in open areas at night than adults (Appendix 3, Table A1). This indicates that TCD in available habitats was lower for adults than for 1^st^ winter birds. For example, in D, 1^st^ winters birds and adults used habitats at night with a mean TCD of 29% and 26%, respectively, while the mean TCD of pseudo-absence within the 3-km buffers was 43% and 36%, respectively.

### Forest habitat selection at the local scale

Selection for local scale habitat characteristics within the forests (from the 402 plots surveyed) was low, but significant for several habitat characteristics (Nagelkerke’s R² = 0.30, Appendix 4, Table A2), particularly during the night (Figure 4, Figure 5, Appendix 4, Table A2). Both habitats availability and selection patterns differed among the three subareas, resulting in distinct local habitat preferences (Figure 4, Figure 5). During the day, woodcocks in A and H preferred plots with higher cover (e.g., in A, the predicted occupancy frequency in plots with 90% cover (75^th^ percentile) was approximately twice that of plots with 60% cover (25^th^ percentile), Figure 4). No significant selection for cover was detected in D, but only 5% of the plots used during the day in D had a cover <60%.

**Figure 4:**
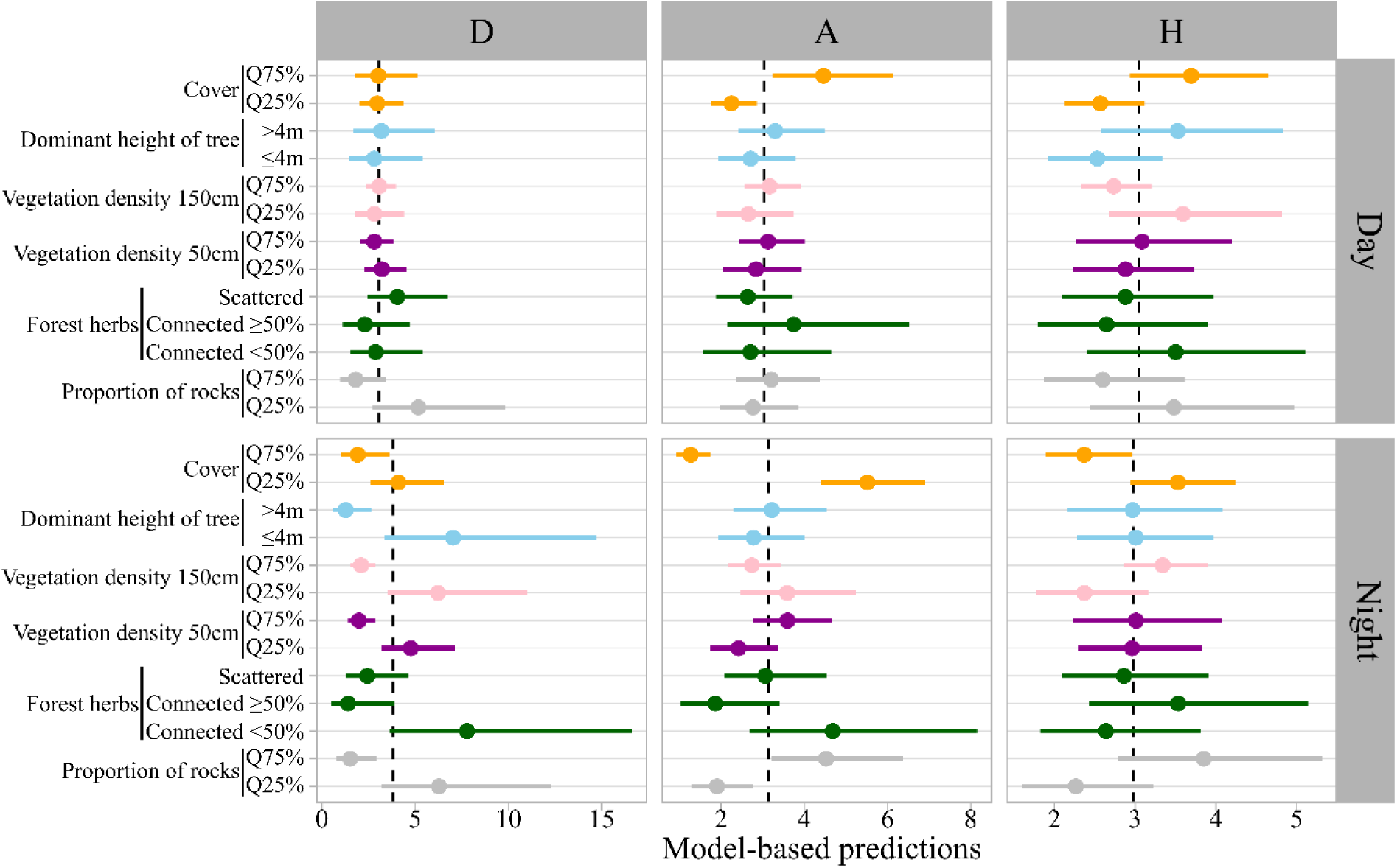
Model based prediction (±95% confidence intervals) of local scale habitat use intensity (number of locations per 100 units as offset) within each subarea. Vertical black dashed lines indicate the average use intensity during day or night period for each subarea. A value higher than this average indicates a rate of use above the average, and vice versa. D: Central Drôme, A: Ardèche gorges, H: Hérault gorges. Q75%: 75^th^ percentile, Q25%: 25^th^ percentile.

**Figure 5:**
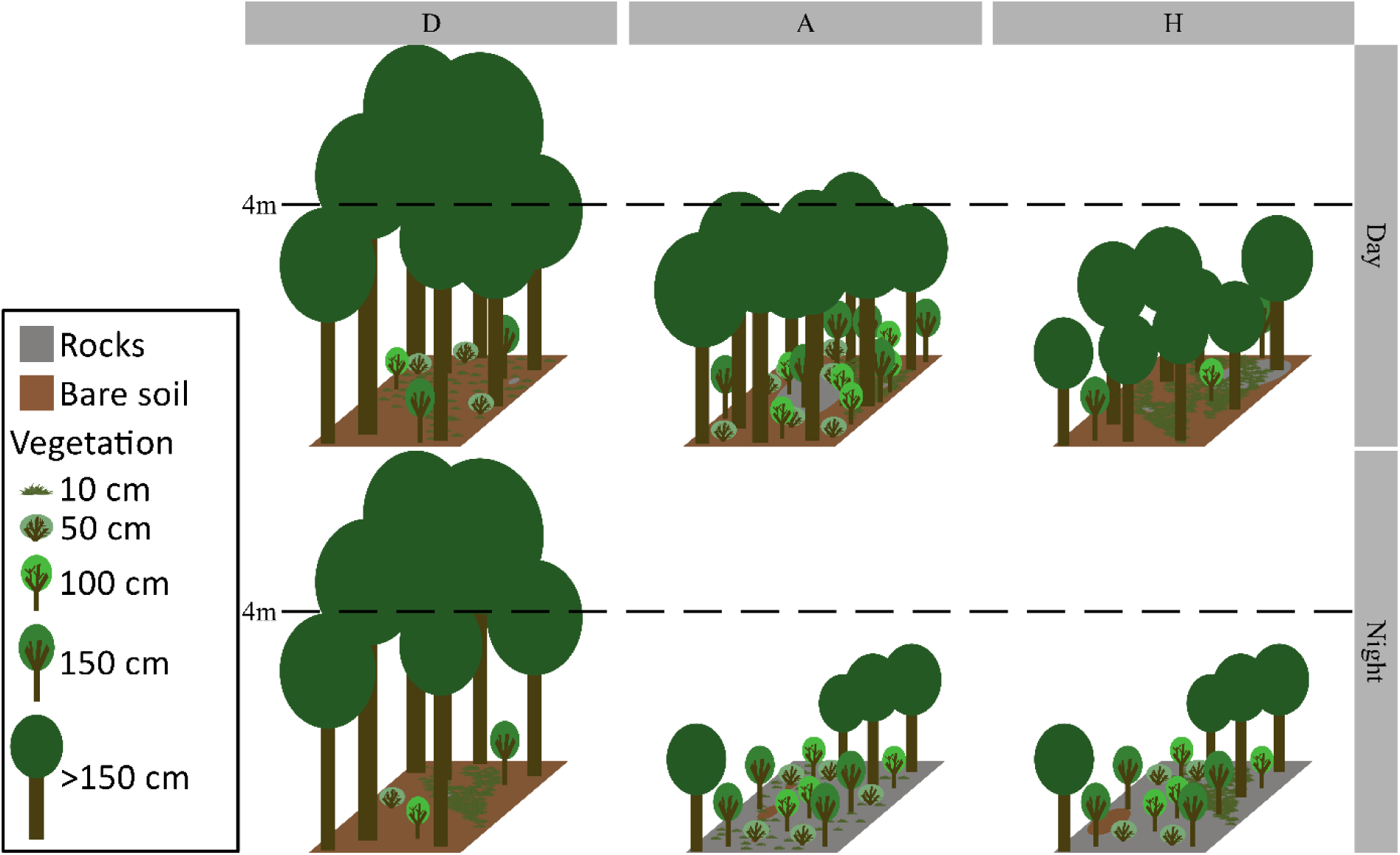
Graphical interpretation of habitats predicted to be more frequently used by woodcocks in each subareas, based on the results of the forest habitat selection model at the local scale. The interpretations were based on a consensus of the 5 plots with the higher predicted value for day and night within the three subareas. The values used for graphical interpretation are available in Appendix 4, Table A3. Subareas: D: Drôme, A: Ardèche gorges, H: Hérault gorges.

During the night, woodcocks in A and H preferentially used plots with lower cover (Figure 4, Figure 5). In addition, woodcocks in D selected plots with a dominant height of trees <4m (Figure 4). Dominant height of trees differed significantly between subareas: trees in A and H were frequently <4m (during the night, 52% of used plots in A and 74% in H) or just above 4m, whereas in D, trees were predominantly well above 4m (during the night, 89% of used plots in D). However, forests with dominant height of trees <4m were rare in D (only 6 plots), and two of these plots accounted for >20% of nocturnal forest locations each for two different individuals. When these two individuals were removed from the analysis, the selection for dominant height of trees <4m in D was no longer significant.

Vegetation density preferences at night also differed among subareas (Figure 4, Figure 5). Woodcocks in D showed a strong preference for lower vegetation density and plots with <50% forest herbs cover, whereas woodcocks in H showed a moderate preference for higher vegetation densities at 150 cm, and no significant preference was detected in A (Figure 4). Finally, rock cover was positively selected at night in A but avoided in D (Figures 4, 5).

## Discussion

Tracking of wintering woodcocks in the Mediterranean region revealed behaviors and habitat selection strategies that contrasted markedly with observations from other parts of their wintering range. This emphasizes the importance of studying species across diverse habitats and under variable climatic conditions to get a complete overview of species-environment relationships.

### Movement patterns & landscape-habitat selection

The most obvious behavioral change concerned the habitat used at night. Typical wintering behavior described for the woodcock consists of using forested habitats during the day and open habitats at night, involving a so-called commuting flight. This behavior is very common in regions with Atlantic climate influence, e.g., occurring 94% of nights in England (Hoodless & Hirons, 2007), and 85% in the north-west part of France (Duriez, Fritz, Said, et al., 2005). Conversely, it is much less frequent in the Mediterranean region. In the northernmost part of the study area, woodcocks commuted to open areas 53% of nights, whereas this occurred in only 11% of nights in the southern part, despite open areas being available within their home range.

Additionally, the analysis of woodcock movements suggested higher diurnal activity, especially in the subarea most strongly influenced by the Mediterranean climate, the Hérault gorges (H). Measuring the activity rate of woodcocks using accelerometry, combined with location data, could provide valuable insights into both their energy expenditure and the timing and locations of feeding under varying climatic conditions (Patterson et al., 2019; Wilson et al., 2006). The activity rate of some woodcocks was recorded during this study and confirmed that the birds in A and H were more active during the day. This aligns with the hypothesis proposed by Duriez, Fritz, Binet, et al. (2005) and with the observations of Hoodless & Hirons (2007), suggesting that woodcocks shift toward diurnal activity under milder climatic conditions. One explanation is that the costs of thermoregulation are lower at milder temperatures, potentially reducing the birds’ overall food requirements and, consequently decreasing the need to forage in riskier open habitats at night (Duriez, Fritz, Binet, et al., 2005). On the other hand, forests may provide shelter from strong winds, which are common in Mediterranean regions and can impose significant thermoregulation costs (Duriez et al., 2004). Indeed, the study region is highly exposed to Mistral, a strong, cold northerly wind that accelerates in the Rhône corridor. However, the effect of wind was not tested in this study, as a proper assessment would require detailed analyses of wind intensity and direction in relation to local topography, vegetation cover, and anthropogenic structures that may act as wind barriers.

### Local habitat selection

When they were in wooded areas at night (47%, 60% and 89% of nocturnal locations in D, A, H, respectively), woodcock selected areas with lower tree cover density in A and H, similar to woodcocks in Central European forests tracked during the breeding period (Sládeček et al., 2023), and with slightly denser vegetation in the shrub stratum in H, but lower in D (Figure 4, Figure 5). Overall, however, the results of habitat selection analyses at a local scale did not detect marked selection for the specific environmental variables measured. Woodcocks used extremely heterogeneous forest habitats depending on the region, with different forest species composition, e.g. mainly tree species of the *Quercus* genus in the studied subareas compared to *Acer pseudoplatanus* and *Fagus sylvatica* in the UK (Hoodless & Hirons, 2007). Counterintuitively, they used habitats with high rock cover at night in A and H, which likely prevents them from probing for earthworms, their principal prey (Granval, 1986), or alternative prey (Aradis et al., 2019). It is unlikely that the small differences in forest habitat selection between day and night were caused by prey availability (Duriez, Ferrand, et al., 2005). To our knowledge, there is no evidence that nocturnal forest habitats characterized by lower tree cover density, higher shrub stratum density, and higher rock cover provided more prey than diurnal habitats. Instead, nocturnal forest habitats were probably selected for protection from predators (Braña et al., 2010). In particular, the tendency of birds from H to use nocturnal habitats with high vegetation density could indicate a prioritization of shelter over terrain traversability at night, which is associated with reduced feeding activity as shown by shorter nocturnal distances of movement patterns in H. Adequate overhead tree and shrub cover may shield woodcocks from nocturnal avian predators, e.g., the Eurasian eagle-owl, *Bubo bubo* (Obuch, 2024), while relatively low tree cover allows for easier take-off in response to ground predators.

### Differences between sex and age classes

Habitat selection and movement patterns were generally consistent between males and females, as well as between adults and 1^st^ winter birds, except for the selection of tree cover density at the landscape scale. During the day, males showed a significantly stronger preference for areas with denser tree cover compared to females. At night, 1^st^ winter birds selected areas with lower tree cover density than adults. 1^st^ winter birds might be compelled to use less favorable wintering areas (e.g., areas with higher predation risk) due to competition or inexperience (Skórka et al., 2016), while the difference between the sexes might be due to small differences in the diet (Aradis et al., 2019). While wintering, there are few dimorphisms or differences in energy requirements between sexes and age classes (Duriez et al., 2004), so few differences in physiological requirements were expected. The sample size for the micro-habitat selection model was probably insufficient to detect such small differences.

### Diet and prey availability

Understanding these behaviors and habitat selection cannot be achieved without better knowledge of diet and prey availability in relation to environmental constraints. The diet in southern subareas relied less on earthworms than in northern areas, showing a higher frequency of millipedes and beetles (Aradis et al., 2019; Granval, 1986), and these prey are not necessarily more abundant in open areas, reducing the incentive for woodcocks to leave the forest. In north-west Europe, earthworms are the main prey items of woodcocks, and they become more available in open field habitats than in woodlands during winter, increasing the incentive for woodcocks to leave the forest (Hoodless & Hirons, 2007). To confirm the effects of prey diversity and distribution on woodcock behavior, it would be interesting to study both the soil macrofauna in the habitats used and to refine our knowledge of woodcock diet in Mediterranean regions (Aradis et al., 2019; Granval, 1986). Assessing diet and prey availability for woodcocks is labor-intensive and requires specialists who can identify taxa at a sufficient taxonomic resolution from different species groups. DNA-based methods could help with species identification, but are not able to provide quantitative information (Gongalsky, 2021; Hoenig et al., 2022; Smith et al., 2008). Future protocols in Mediterranean regions should include the identification of additional prey species (e.g., Coleoptera and Diplopoda, Aradis et al., 2019) to achieve greater concordance with woodcock prey in Mediterranean regions. In addition, a significant number of habitats should be included in food resource studies to account for differences in the habitat selection strategies of woodcocks wintering in different climatic regions.

### Effect of environmental conditions on behavior and population management

Temperature and soil humidity affected movement patterns and habitat selection, reinforcing the idea that prey availability played a key role in the observed behaviors. Specifically, woodcocks increase the frequency of commuting flights to open habitats at night when the soil became wetter and temperatures were milder, the latter effect being even stronger in the subareas most influenced by the Mediterranean climate (A and H). They also selected nocturnal habitats with higher tree cover density at landscape scale when the soil became drier and temperatures higher. They appeared to change their diel activity pattern, e.g. reducing diurnal activity when temperatures were higher (as shown by reduced diurnal distances), although further analyses of activity rates and associated behavior are needed to confirm this change. Forest habitats can help maintain soil moisture during drier periods in Mediterranean regions (Sardans & Peñuelas, 2013), ensuring an adequate food supply in forests.

In addition, we observed that the proportion of 1^st^ winter birds among the birds hunted or trapped was much lower in dry winters in the southern subarea (*Réseau Bécasse - Lettre d’information n° 27*, 2018), whereas this proportion was much higher in northern parts of France. This suggests that inexperienced 1^st^ winter birds tend to avoid these areas when dry, while adult birds are more likely to return and remain even in less suitable environmental conditions. Furthermore, we recorded eight woodcocks relocating from the southern subarea (H) during winter 2022, while woodcocks are known to be very faithful to their wintering areas throughout the winter (Ferrand et al., 2008), suggesting that the wintering area no longer provided the necessary conditions to meet the birds needs, probably due to a lack of food availability. One of the eight individuals relocated was recaptured 100 km away and was in poor body condition (260 g compared to 320 g when first captured). Teitelbaum et al. (2023) identified 92 migratory bird species that made post-migration movements during the non-breeding season, indicating that such movement patterns are probably not rare events. However, the fact that most of the tagged birds in the southern subarea H (30/38 woodcocks) preferred to stay in a known location during dry winters rather than relocate suggests that relocation comes at a significant cost compared to maintaining site fidelity for the woodcock. This behavior of relocating during winter could serve as an early warning signal of declining habitat suitability for wintering woodcocks in Mediterranean regions. Further research is needed to identify the factors that trigger these relocations. Considering woodcocks, we hypothesize that food availability is the most important factor (Duriez, Ferrand, et al., 2005; Martín et al., 2023). In addition, the relocation of wintering woodcocks may lead to changes in local population density and could indicate vulnerability of the species which would require a revision of population management. Similar adaptations of population management already exist during prolonged cold spells; woodcock hunting is suspended and enhanced monitoring of the species is implemented (population inventories and assessment of the physiological condition, mortality and behavior).

### Implications of behavioral plasticity for habitat management

The differences in movement and habitat selection strategies among the three Mediterranean subareas probably reflected differences in climate influence, as well as the structure and distribution of forests and open habitats, all of which affect the diversity and density of prey and predators. Forests in the southernmost subareas were used for both shelter and foraging, whereas woodcocks in the northern subarea more often segregated feeding in open areas and resting in forests. Such behavioral plasticity has already been observed in several species living in different habitats or under changing conditions caused by human activities or climatic events (e.g., foraging plasticity, Gilmour et al., 2018; migratory decisions, Schumm et al., 2022; or changes in home range size after wildfire, Kreling et al., 2021). However, this plasticity may be limited in some contexts, and its effects on individual fitness are often unknown. The relocation behavior observed for 8 birds in 2022 was a clear indication that individuals adaptive abilities remain limited. Therefore, habitat conservation and management remain crucial to maintaining the species throughout its wintering range and must account for variability in wintering behavior between geographical areas. For example, in the Drôme (D), maintaining open habitats with high earthworm densities could be beneficial, whereas in the Ardèche (A) and the Hérault (H) gorges, a mosaic of dense and sparse forests could be more advantageous than maintaining open areas. Overall, our results reaffirm the importance of preserving heterogeneous habitats that provide multiple options for species to persist locally under varying conditions (Oliver et al., 2010; Suggitt et al., 2018). Even within a single climatic region (<200 km apart), woodcock ecology varies considerably, highlighting the complexity of defining suitable habitat for a species.

Although woodcocks are widespread and a well-studied species, our study reveals important knowledge gaps in their ecology. This highlights the misunderstanding we often have about a variety of species because they are only studied in a small portion of their temporal and geographic range, even though animal responses to available habitat are often context-dependent (Broekman et al., 2022; Constible et al., 2010). Our results therefore highlight a well-known but often overlooked principle in species-habitat relationships: extrapolating results beyond the geographic or temporal range of the studied population is precarious. To refine the assessment of habitat suitability, it is essential to study a sufficiently representative variety of environments within the distribution and behavior-specific habitat of a species (Roever et al., 2014).

## Conclusion & perspectives

The study of woodcocks wintering in the Mediterranean region shows the remarkable behavioral flexibility of this species at the climatic edges of its range. However, this flexibility complicated predictions of the woodcocks’ geographic range dynamics under climate change (Oldfather et al., 2020). While behavioral adaptations appear to mitigate some challenges, their long-term effectiveness remains unclear. Our results suggest that behavioral adaptations, such as changes in habitat use, are often preferred over relocation to more suitable wintering areas. Nevertheless, we lack data to determine whether these behavioral changes maintain or worsen fitness benefits. Measuring fitness is crucial to assess whether the persistence of woodcocks in Mediterranean regions during wintering is at risk in the near future.

The Mediterranean region is probably one of the most vulnerable areas to exceeding the climatic conditions tolerable by woodcocks due to ongoing climate change. The effects of climate change in the Mediterranean region, including decreasing precipitation combined with an increase in the frequency and severity of droughts, could significantly impact local fauna, but are not yet well understood (Cramer et al., 2018; Ferreira et al., 2022) and present a pressing concern (Moatti & Thiébault, 2016). For instance, drought conditions can induce physiological stress (Dezetter et al., 2022), alter prey communities (Martín et al., 2023) and influence habitat use (Pearse et al., 2024). This underlines the urgent need to study species-specific ecological requirements in Mediterranean ecosystems to predict how climate change might affect species distribution and survival and to develop effective conservation strategies.

## Appendices

Supplementary data to this article can be found online in Zenodo (https://doi.org/10.5281/zenodo.18605303)

## Acknowledgement

We warmly thank the professionals, students and volunteers for their assistance in the data collection: Luc Aurousseau, Christophe Mathez, Guillaume Nard, Damien Dubosc, Ludovic Fallais, Sébastien Blanchard, Jean-Claude Etienne, Maxime Passerault and Damien Coreau. We are also grateful to Elodie Coudour for her valuable support with administrative tasks. This project was funded by the ecocontribution (Office Français de la Biodiversité, grant no. 1251).

## Notes

https://doi.org/10.5281/zenodo.18605303

### Competing Interest Statement

The authors have declared no competing interest.

### Summary of Updates

add the PCI Zoology badge and remove line numbering.

